# Graminoids vary in functional traits, carbon dioxide and methane fluxes in a restored peatland: implications for modeling carbon storage

**DOI:** 10.1101/2021.05.27.445980

**Authors:** Ellie M. Goud, Sabrina Touchette, Ian B. Strachan, Maria Strack

## Abstract

One metric of peatland restoration success is the re-establishment of a carbon sink, yet considerable uncertainty remains around the timescale of carbon sink trajectories. Conditions post-restoration may promote the establishment of vascular plants such as graminoids, often at greater density than would be found in undisturbed peatlands, with consequences for carbon storage. Although graminoid species are often considered as a single plant functional type (PFT) in land-atmosphere models, our understanding of functional variation among graminoid species is limited, particularly in a restoration context. We used a traits-based approach to evaluate graminoid functional variation and to assess whether different graminoid species should be considered a single PFT or multiple types. We tested hypotheses that greenhouse gas fluxes (CO_2_, CH_4_) would vary due to differences in plant traits among five graminoid species in a restored peatland in central Alberta, Canada. We further hypothesized that species would form two functionally distinct groupings based on taxonomy (grass, sedge). Differences in gas fluxes among species were primarily driven by variation in leaf physiology related to photosynthetic efficiency and resource-use, and secondarily by plant size. Multivariate analyses did not reveal distinct functional groupings based on taxonomy or environmental preferences. Rather, we identified functional groups defined by continuous plant traits and carbon fluxes that are consistent with ecological strategies related to differences in growth rate, resource-acquisition, and leaf economics. These functional groups displayed larger carbon storage potential than currently-applied graminoid PFTs. Existing PFT designations in peatland models may be more appropriate for pristine or high-latitude systems than those under restoration. Although replacing PFTs with continuous plant traits remains a challenge in peatlands, traits related to leaf physiology and growth rate strategies offer a promising avenue for future applications.

## Introduction

Peatlands are globally important carbon sinks, storing approximately one-third of the world’s terrestrial soil carbon with an estimated stock over 600 Gt (Loisel et al., 2021; Loisel et al., 2014; Yu et al., 2010). In Canada, peatlands cover an estimated 120 million ha of land surface, the second largest cover of peatlands in the world (Vitt, 2013). Canada is a major producer and exporter of peat for horticultural uses, producing about 1.3 million metric tons of peat per year. Approximately 34,000 ha of peatlands have been drained for peat harvesting, with approximately 58% actively in use (Environment Canada 2020). Considerable carbon is released during the harvesting process primarily in the form of greenhouse gases carbon dioxide (CO_2_) and methane (CH_4_). Post-harvested peatlands may remain a net source of carbon in the absence of restoration efforts (Cleary et al., 2005; Nugent et al., 2019). To offset the carbon cost of peat harvesting and help mitigate potential feedbacks to climate change, post-harvested peatlands are restored with the goal of returning them to a persistent carbon sink (Chimner et al., 2017; González & Rochefort, 2014).

Despite notable successes, considerable uncertainty remains around post-restoration carbon flux trajectories, in part due to a limited number of long-term studies (González & Rochefort, 2014). To complement long-term studies, process-based models are used to describe the environmental controls on carbon dynamics and predict future carbon accumulation (Frolking et al., 2010). A key component of land-atmosphere models is vegetation, as plants directly control carbon cycling through photosynthesis, respiration, decomposition, and CH_4_ release. To simplify model structure, vegetation are represented as ‘plant functional types’ (PFTs), with different species assigned to a PFT based on assumed similarity in function, environmental tolerance, or contribution to peat growth (Frolking et al., 2010; Heinemeyer et al., 2010; Wu & Blodau, 2013).

While the goal of PFT is to represent species with similar function in a modeling context, generic PFT labels such as ‘moss’, ‘shrub’ and ‘graminoid’ are now commonly used in the peatland ecological literature, often forming the basis of experimental treatments (Goud et al., 2017; Robroek et al., 2016; Rupp et al., 2019). While a range of species are assigned to a single PFT, current peatland PFT designations are based on only a few studies predominantly from pristine, high-latitude peatlands (e.g., Frolking et al. 2010; Laine et al. 2012; Tuittila et al. 2012). Given that different species within a PFT may vary substantially in carbon accumulation and greenhouse gas emissions based on local environmental conditions and disturbances (Goud et al., 2017; Lai et al., 2014), some PFT designations may not be relevant for managed and restored peatlands.

In a restoration context, differences in post-disturbance conditions (e.g., soil compaction and moisture availability) substantially alter plant community composition relative to undisturbed sites, especially in the first few years, in large part because of the early establishment of graminoid species (González & Rochefort, 2019; Graf et al., 2008). Graminoids are grass-like vascular plants, including grasses (family Poaceae), sedges (Cyperaceae), rushes (Juncaceae), arrow-grasses (Juncaginaceae), and quillworts (Isoetes). Graminoids often have much larger rates of CO_2_ and especially CH_4_ release relative to other peatland plants (Goud et al., 2018; Lai et al., 2014; Maria Strack et al., 2017). Consequently, restored peatlands with considerable graminoid cover have different rates of carbon storage compared to natural counterparts (Strack et al., 2016). Specifically, due to the greater radiative forcing of CH_4_ over CO_2_, extensive graminoid cover that potentially suppresses subsequent peatland successional stages could compromise a restored peatland’s ability to be a net carbon sink over time (Frolking et al., 2006; Strack et al., 2006). As such, there is considerable interest in accurately modeling carbon exchange in graminoid-dominated peatlands.

Despite the importance of graminoids to peatland carbon cycling, especially for restored peatlands, our understanding of graminoid functional variation is limited. While rates of CO_2_ and CH_4_ fluxes vary in predictable ways with hydrology and peat temperature (Lai et al., 2014; Strack et al., 2016), biotic drivers of carbon cycling within graminoids are not well established. This knowledge gap is further reflected in the inconsistent representation of graminoids in current ecological studies and land-atmosphere models, where all graminoids are either categorized as a single PFT (Chaudhary et al., 2017; Wania et al., 2009; Wu & Blodau, 2013) or subdivided based on taxonomy (i.e., ‘grass’, ‘sedge’, ‘rush’) (Frolking et al., 2010; Heinemeyer et al., 2010) or environmental preferences (i.e., ‘minerotrophic’, ‘ombrotrophic’) (Frolking et al., 2010; Quillet et al., 2015; Tuittila et al., 2012). In other graminoid-dominated ecosystems such as grasslands and marshes, variation in carbon storage has been linked to plant traits such as height (Klumpp & Soussana, 2009; Radabaugh et al., 2017) and leaf nitrogen content (Long et al., 2019; Tripathee & Schäfer, 2015). Although there are fewer trait-based studies in peatlands, recent work demonstrates carbon cycling among divergent plant groups (i.e., moss, shrub, graminoid) can be predicted by functional traits such as leaf area (Goud et al., 2017; Korrensalo et al., 2016) and nitrogen content (Girard et al., 2020). Applying plant functional traits to identify mechanistic drivers of greenhouse gas fluxes within graminoids has the potential to advance our understanding of post-restoration ecological dynamics in general, and to better represent graminoid contributions to carbon fluxes in land-atmosphere models (VanBodegom et al., 2011).

Here, we used a traits-based approach to evaluate graminoid functional variation in a restored peatland in central Alberta, Canada. Our objectives were two-fold: 1) determine functional differences among graminoid species to better understand anatomical and physiological mechanisms underlying variation in CO_2_ and CH_4_ fluxes; 2) assess whether graminoid species should be considered a single PFT or multiple based on taxonomy or environmental preferences. We hypothesized that CO_2_ and CH_4_ fluxes would vary among species due to differences in traits related to plant size and leaf physiology. We further hypothesized that species’ functional groupings would be more related to taxonomy than environmental preferences. To test these hypotheses, we measured gas fluxes and six functional traits in five dominant graminoid species that vary in morphology, taxonomy, and environmental preferences. We measured plant traits used in PFT classifications (i.e., above-ground biomass, leaf nitrogen content, photosynthetic quantum yield) and traits that relate to plant size and resource acquisition (i.e., plant height, leaf carbon and nitrogen stable isotope ratios).

## Methods

### Study site and experimental setup

Measurements were made in a restored ombrotrophic bog located 17 km southeast of Entwistle, Alberta, Canada (53°27’26”N, 114°53’04”W). The climate is cool-continental, with a normal (1981-2010) mean annual air temperature of 3.5 °C and total annual precipitation of 551 mm. Peak growing season (June-August) mean temperature and total precipitation are 15.5 °C and 225 mm, respectively (Environment Canada 2020). Peat was harvested from 2000 - 2012 and subsequently restored using the moss-layer transfer technique (González & Rochefort 2014) in the autumn and winter of 2012. We focused our measurements on five target species: American Slough Grass (*Beckmannia syzigachne* (Steud.) Fernald), Bluejoint (*Calamagrostis canadensis* (Michx.) P. Beauv.), Silvery Sedge (*Carex canescens* L.), Tussock Cottongrass (*Eriophorum vaginatum* L.), and Woolgrass (*Scirpus cyperinus* (L.) Kunth). All are obligate wetland species with C3 metabolism and aerenchyma tissue (Ball et al., 2002; Clark & Kellogg, 2007). *Beckmannia* and *Calamagrostis* are grasses (Poaceae) while *Carex*, *Eriophorum* and *Scirpus* are sedges (Cyperaceae). *Beckmannia syzigachne* is an annual, the other four are perennials. *Eriophorum vaginatum* is typically associated with ombrotrophic wetlands (e.g., bogs), while the remaining four species occur in minerotrophic wetlands (e.g., fens, marshes, swamps) (Gleason & Cronquist, 1991). In May 2016, we established 20 square plots (0.60 m x 0.60 cm) for gas flux measurements centered around a target species (n = 4 plots per species). Grooved aluminum collars were permanently installed to allow for repeated gas flux measurements in the same location to minimize disturbance to the peat and vegetation. We additionally established 50 square plots (0.25 m x 0.25 m) for destructive plant sampling centered around a target species (n = 10 plots per species).

### Carbon dioxide and methane flux measurements

Gas fluxes were measured at the whole-plant level using chambers, which included multiple plant individuals and soil heterotrophic respiration and CH_4_ fluxes. We measured net ecosystem CO_2_ production (NEP, μmol CO_2_ m^-2^ s^-1^), ecosystem respiration (ER, μmol CO_2_ m^-2^ s^-1^), gross primary productivity (GPP, μmol CO_2_ m^-2^ s^-1^) and CH_4_ (mg m^-2^ d^-1^) fluxes every second week from May to September 2016. To measure NEP, we used the closed dynamic chamber method (Alm et al., 2007) using a transparent acrylic chamber (60 cm × 60 cm × 30 cm) equipped with a cooling system. Two battery-operated fans circulated headspace air within the chamber, blowing past a copper coil containing cold water circulating from a cooler with ice to ensure minimal heating during the measurements. For plants taller than 60 cm, an acrylic extension of 60 cm high with two additional battery-operated fans was added under the chamber to prevent damage to the stem of the plants. A thermocouple thermometer and a photosynthetically active radiation (PAR, μmol quanta m^-2^ s^-1^) sensor recorded environmental conditions within the chamber during measurements. CO_2_ concentrations were determined using a portable infrared gas analyzer (IRGA; PPsystems EGM-4, Massachusetts, USA) and the change in CO_2_ concentration over a two-minute period was determined *in situ* every 15 seconds. The linear rate of CO_2_ concentration increase was used to calculate the flux. To achieve various PAR levels for estimating light response curves, we measured ambient light conditions at the time of measurement and used mesh covers to partially shade the chambers. We measured ER by covering the chamber with an opaque tarp to achieve PAR = 0. The difference between NEP and ER was used to calculate GPP. We adopt the ecological sign convention that positive NEP values indicate uptake of CO_2_ by the ecosystem while negative values indicate a release of CO_2_ to the atmosphere.

We measured CH_4_ fluxes using the closed static chamber method (Alm et al., 2007) with opaque chambers (60 cm × 60 cm × 30 cm) equipped with fans for air circulation, a thermocouple, and an acrylic extension of 60 cm high with two additional battery-operated fans to accommodate plants taller than 60 cm. Chambers were placed on sampling collars for 35 minutes, and four air samples were taken at 5, 15, 25, and 35 minutes into a 20 mL syringe and then transferred to 12 mL pre-evacuated vials (Exetainers, Labco Ltd., UK) to measure the change in concentration of CH_4_ inside the chamber head space. CH_4_ concentrations were determined in the laboratory with a gas chromatograph (GC-2014 Gas Chromatograph, Shimadzu Scientific Instruments, Kyoto, Japan) with a flame ionization detector. The CH_4_ flux was determined as the slope of the linear change in concentration versus time over the 35-minute sampling period. Before determining the CH_4_ flux, data was quality controlled where measurements with concentrations at 5 minutes that were higher than 5 ppm followed by a decline in concentration, or erratic concentration changes likely associated with ebullition events were removed from the data set. Cases where concentration was less than 5 ppm and did not change more than the precision of the gas chromatograph (10%) were assigned a flux value of zero; zero fluxes were retained.

Fluxes without a significant regression correlation coefficient (R^2^ > 0.90 at *p* < 0.05) were rejected, with a rejection rate of < 3% and < 9% for CO_2_ and CH_4_, respectively. To obtain peak growing season fluxes, we averaged CO_2_ and CH_4_ flux data from June - July. We retained NEP and GPP data during the peak growing season from measurements at PAR > 1000 μmol m^-2^ s^-1^ for inter-specific comparisons. With each flux measurement, water table depth below the peat surface was measured from a standpipe next to the collars, and soil temperature (°C) was measured at 2, 5, 10, 15 and 20 cm below the peat surface with a portable thermocouple probe (Digi-Sense Type-K, Oakton Instruments, IL, USA).

### Vegetation sampling and plant traits

In August 2016, we measured maximum plant height (m) in all plots (n = 14 per species). In each destructive-sampling plot, aboveground plant material was clipped and separated between living biomass of the target species and other non-target plant material. Target species biomass was further separated into stem and leaf material. Leaf and stem samples were oven-dried for 48 hours at 60 °C in a mechanical convection oven (Heratherm OMS100, Thermo Scientific, Massachusetts, USA).

Leaf nitrogen percent element (N) and carbon and nitrogen stable isotope ratios (δ^13^C, δ^15^N) were measured using a continuous flow isotope ratio mass spectrometer (Thermo Scientific Delta Plus XL) coupled to an elemental analyzer (Costech ECS 4010). Isotope ratios are expressed as δ values (per mil):

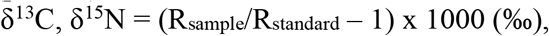

where R_sample_ and R_standard_ are the ratios of heavy to light isotope of the sample relative to the international standards for C and N, Vienna-Pee-Dee Belemnite and atmospheric nitrogen gas (N_2_), respectively. Samples were analysed at the University of Waterloo Environmental Isotope Laboratory, Waterloo, ON.

To calculate plot-level quantum efficiency (φ), we fitted light response curves to NEP data obtained across all PAR value for each species using the ‘light.response’ function in the R Bigleaf package (Knauer et al., 2018). The curve is described by a rectangular hyperbola:

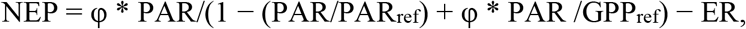

where NEP is net ecosystem production, φ is the initial slope of the curve (quantum efficiency, μmol CO_2_ m^-2^ s^-1^)/(μmol quanta m^-2^ s^-1^), GPP^ref^ is the GPP at a reference light-saturating PAR of 2000 μmol quanta m^-2^ s^-1^, and ER is dark respiration.

### Statistical analyses

We compared variation among variables using one-way analysis of variance (ANOVA) with repeated measures for gas fluxes and without repeated measures for plant traits. We calculated Pearson correlation coefficients among NEP, GPP, ER and CH_4_ (log-transformed) with above-ground biomass, plant height, leaf carbon and nitrogen stable isotope compositions (δ^13^C, δ^15^N), leaf N content, and leaf quantum efficiency (φ). For the plots without flux data (i.e., destructive sampling plots), we used the species’ means values calculated from the gas flux plots. We used cluster analysis and principal components analysis to identify functional groups based on variation in species’ descriptors (i.e., traits, average peak growing season CO_2_ and CH_4_ fluxes). We performed Ward clustering using the ‘agnes’ function in the ‘cluster’ R package on a distance matrix computed from a Pearson correlation matrix among descriptors (Maechler et al., 2017). Ward clustering is a hierarchical agglomerative clustering method that considers all species as being initially separate from each other, and proceeds by successively grouping species into larger and larger clusters until they are all are encompassed into a single cluster (Legendre & Legendre, 2012). We used concordance analysis based on Kendall’s coefficient of concordance (*W*) to identify how many clusters are, in fact, significantly distinct and which plots are significantly contributing to each cluster using the function ‘kendall.global’ in the ‘vegan’ R package, with 999 permutations. Then, *a posteriori* tests of the contribution of individual plots to the concordance of their cluster were computed using the function ‘kendall.post’. We performed principal components analysis using the ‘rda’ function in the ‘vegan’ R package (Oksanen et al., 2019) to identify species assemblages and to compare PCA groupings with those identified from the cluster analysis. PCA preserves the Euclidean distance among descriptors and assumes linear relationships, allowing us to further assess the contribution of descriptors to variation among species.

## Results

### Site conditions

Climate during 2016 was comparable to long-term normals, with a mean annual temperature of 5.5 °C and total annual precipitation of 523.5 mm. Peak growing season (June-August) conditions were similar in mean temperature (16 °C) and received slightly more precipitation (278 mm) compared to the long-term normal (225 mm). Peat water table position and temperature from 2-20 cm did not vary among plots (all p > 0.05).

### Variation in gas fluxes and plant traits among species

Species varied in NEP (F = 26.6, p < 0.0001), GPP (F = 15.7, p < 0.0001), ER (F = 6.4, p = 0.0002), and CH_4_ fluxes (F = 9.1, p < 0.0001). *Scirpus* had the largest NEP, GPP and CH_4_ fluxes, followed by *Eriophorum* (Figure 1 a-b, d). *Eriophorum* had the largest ER (Figure 1 c). *Beckmannia*, *Calamagrostis*, and *Carex* had similar NEP, GPP, ER and CH_4_ fluxes (Figure 1 a-d). Plant traits also varied among species. *Scirpus* had the largest above-ground biomass, followed by *Eriophorum. Beckmannia*, *Calamagrostis*, and *Carex* had similar biomass (Figure 2 a). *Scirpus* was also the tallest species, followed by *Beckmannia. Calamagrostis*, *Carex*, and *Eriophorum* that were of similar height (Figure 2 b). *Scirpus* was the most light-use efficient (largest φ), followed by *Eriophorum* (Figure 2 c). *Beckmannia* had the largest leaf N, *Calamagrostis* had the smallest, while *Carex, Scirpus* and *Eriophorum* had similar, intermediate leaf N (Figure 1 d). *Calamagrostis* had the most negative (relatively depleted) δ^13^C, while *Carex* had the least negative (relatively enriched) δ^13^C (Figure 2 e). *Beckmannia, Calamagrostis*, and *Carex* had the most positive (relatively enriched) δ^15^N, while *Scirpus* and *Eriophorum* had the most negative (relatively depleted) δ^15^N (Figure 2 f).

**Figure 1:**
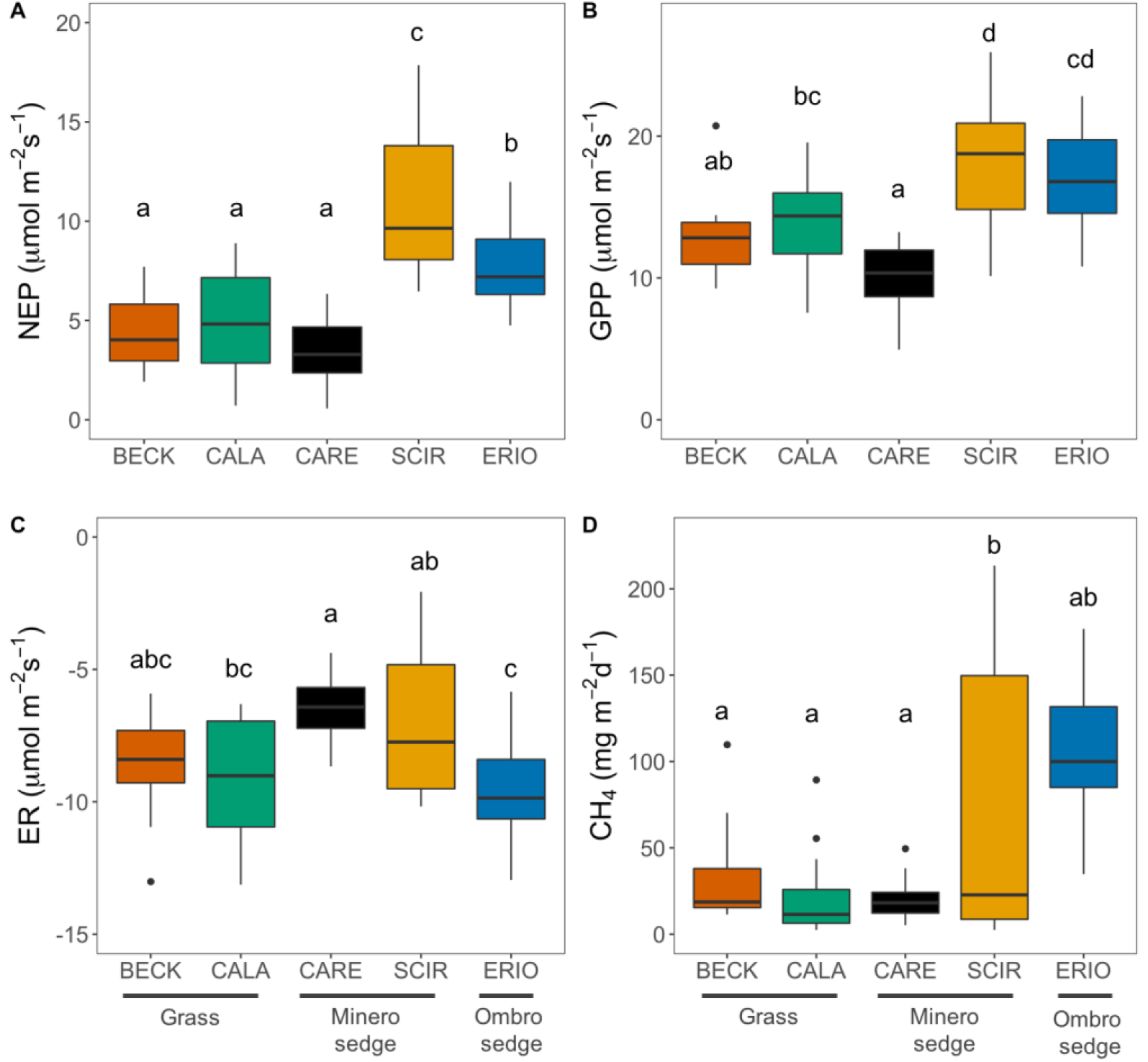
Variation in peak growing season (a) net ecosystem CO_2_ production (NEP), (b) gross primary productivity (GPP), (c) ecosystem respiration (ER), and (d) methane emissions (CH_4_) among five graminoid species in a restored peatland. BECK = *Beckmannia syzigachne*, CALA = *Calamagrostis canadensis*, CARE = *Carex canescens*, SCIR = *Scirpus cyperinus*, ERIO = *Eriophorum vaginatum*. ‘Grass’ refers to species in the Poaceae family, ‘Minero sedge’ and ‘Ombro sedge’ refers to species in the Cyperaceae family from minerotrophic and ombrotrophic environments, respectively. Species that share the same letters are statistically indistinguishable based on Tukey post-hoc tests. Data are from plot means (n = 14 per species).

**Figure 2:**
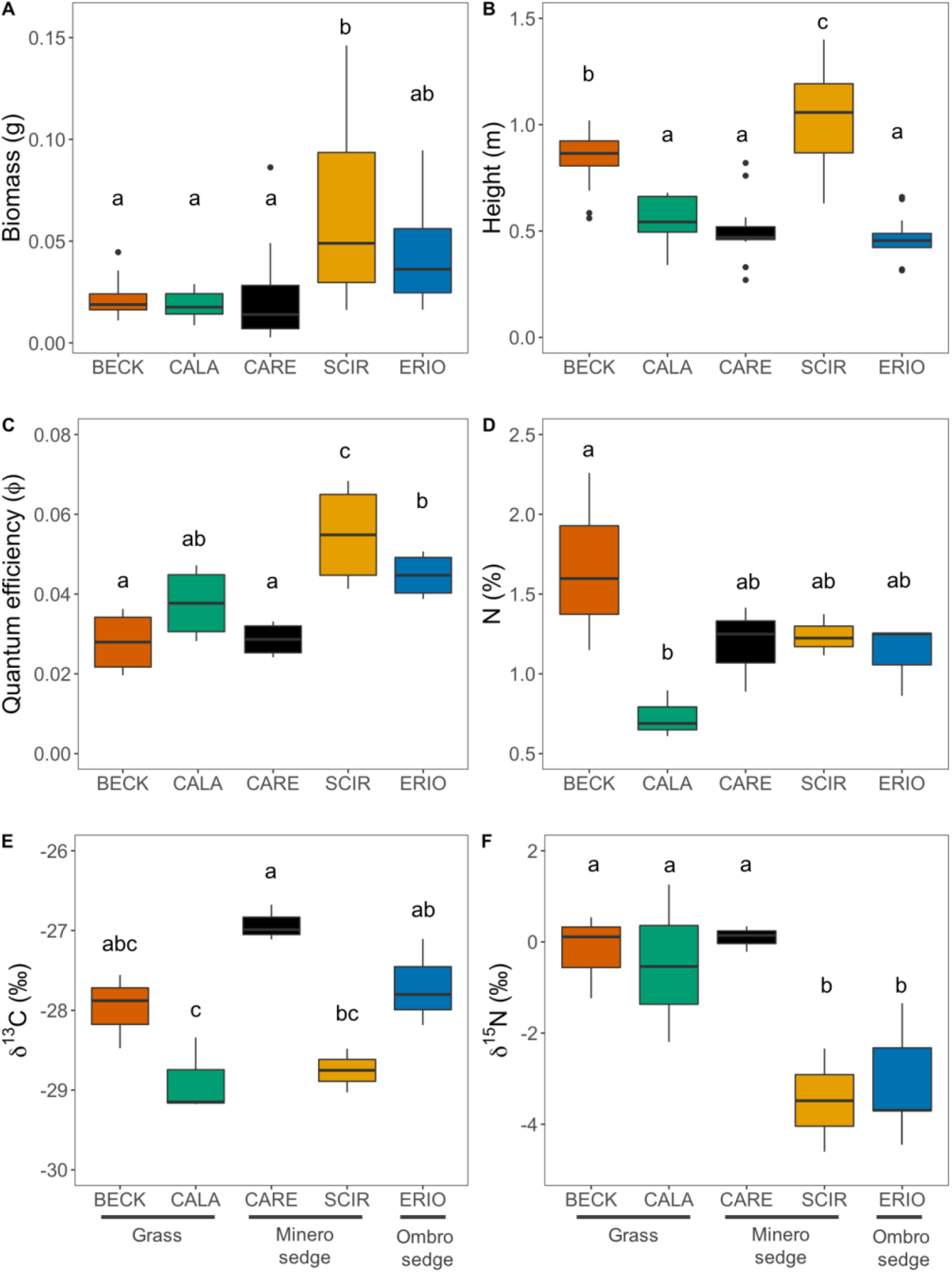
Variation in (a) above-ground biomass (g), (b) plant height (m), (c) quantum efficiency (φ), (d) leaf nitrogen (N,), (e) leaf carbon stable isotope ratio (δ^13^C), and (f) leaf nitrogen stable isotope ratio (δ^15^N) among five graminoid species in a restored peatland. BECK = *Beckmannia syzigachne*, CALA = *Calamagrostis canadensis*, CARE = *Carex canescens*, SCIR = *Scirpus cyperinus*, ERIO = *Eriophorum vaginatum*. ‘Grass’ refers to species in the Poaceae family, ‘Minero sedge’ and ‘Ombro sedge’ refers to species in the Cyperaceae family from minerotrophic and ombrotrophic environments, respectively. Species that share the same letters are statistically indistinguishable based on Tukey post-hoc tests. Data are from plot means (n = 14 per species).

### Plant anatomical and physiological drivers of gas fluxes

Biomass and φ positively correlated with NEP, GPP and CH_4_ (Table 1). Height positively correlated with NEP, GPP and ER. δ^13^C negatively correlated with NEP and GEP, and positively correlated with ER. δ^15^N negatively correlated with NEP, GPP, and CH_4_. Leaf N did not correlate with any gas fluxes. The strongest predictors of NEP and GPP were δ^15^N and φ, followed by δ^13^C, biomass, and plant height. The strongest predictors of ER were δ^13^C and plant height. δ^15^N was the strongest predictor of CH_4_, followed by φ and biomass (Table 1).

**Table 1:**
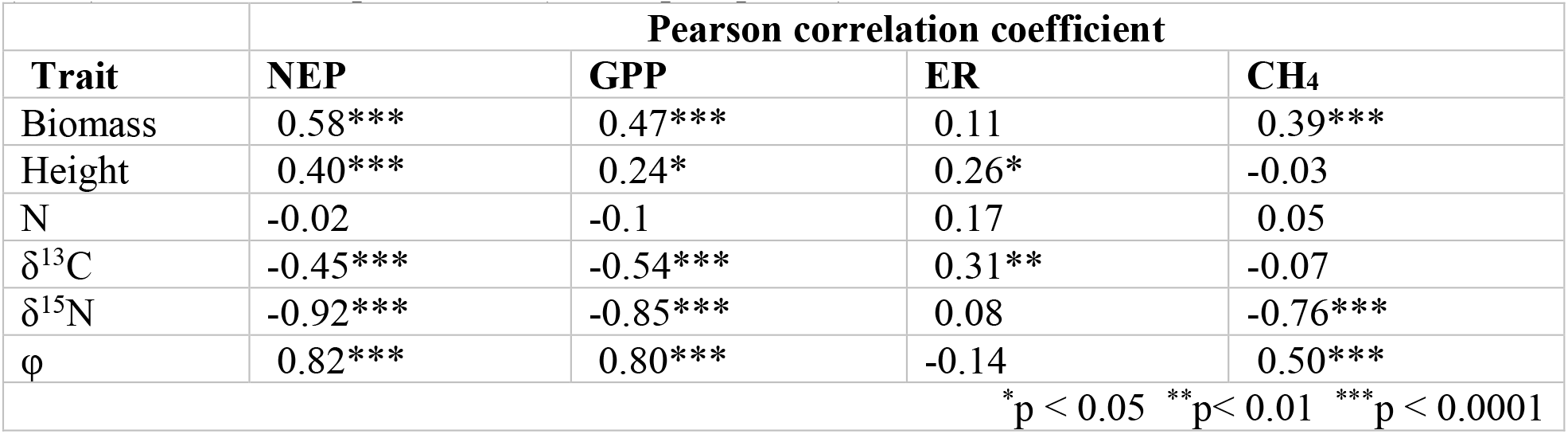
Pearson correlation coefficients between gas fluxes and plant traits among five graminoid species in a restored peatland. Dependent variables are net ecosystem CO_2_ production (NEP), gross primary productivity (GPP), ecosystem respiration (ER), and methane (CH_4_, log-transformed) fluxes. Independent variables are above-ground biomass, plant height, quantum efficiency (φ), leaf carbon isotope composition (δ^13^C), and leaf nitrogen isotope composition (δ^15^N). Data are from plot means (n = 14 per species).

### Graminoid functional groupings

Cluster analysis segregated the 70 plots into five distinct clusters that each contained a mixture of 2-3 different species. Three clusters contained all *Scirpus* and *Eriophorum* plots plus a single *Beckmannia* and *Carex*, and two clusters contained only *Beckmannia*, *Calamagrostis* and *Carex* (Figure 3 a). All five clusters and their associated plots were significant based on Global Kendall tests (0.44 < W < 0.54, p < 0.001) and *a posteriori* Kendall tests (0.97 < W < 0.99, p < 0.001). Plots were distributed along three PC axes that together explained 81% of the variation. The first PC axis (50%) was primarily associated with variation in NEP, GPP, δ^15^N, and φ. The second PC axis (18%) was primarily associated with variation in leaf N, plant height, ER, and biomass. The third PC axis (13%) was primarily associated with variation in δ^13^C and CH_4_ (Table 2). As in the cluster analysis, *Beckmannia* and *Carex* completely overlapped in the PCA biplot, and *Calamagrostis* overlapped with them along PC2. *Scirpus* and *Eriophorum* overlapped along PC1 and PC2. *Scirpus*, *Calamagrostis* and *Eriophorum* were somewhat more differentiated along PC3 (Figure 3 b-c).

**Figure 3:**
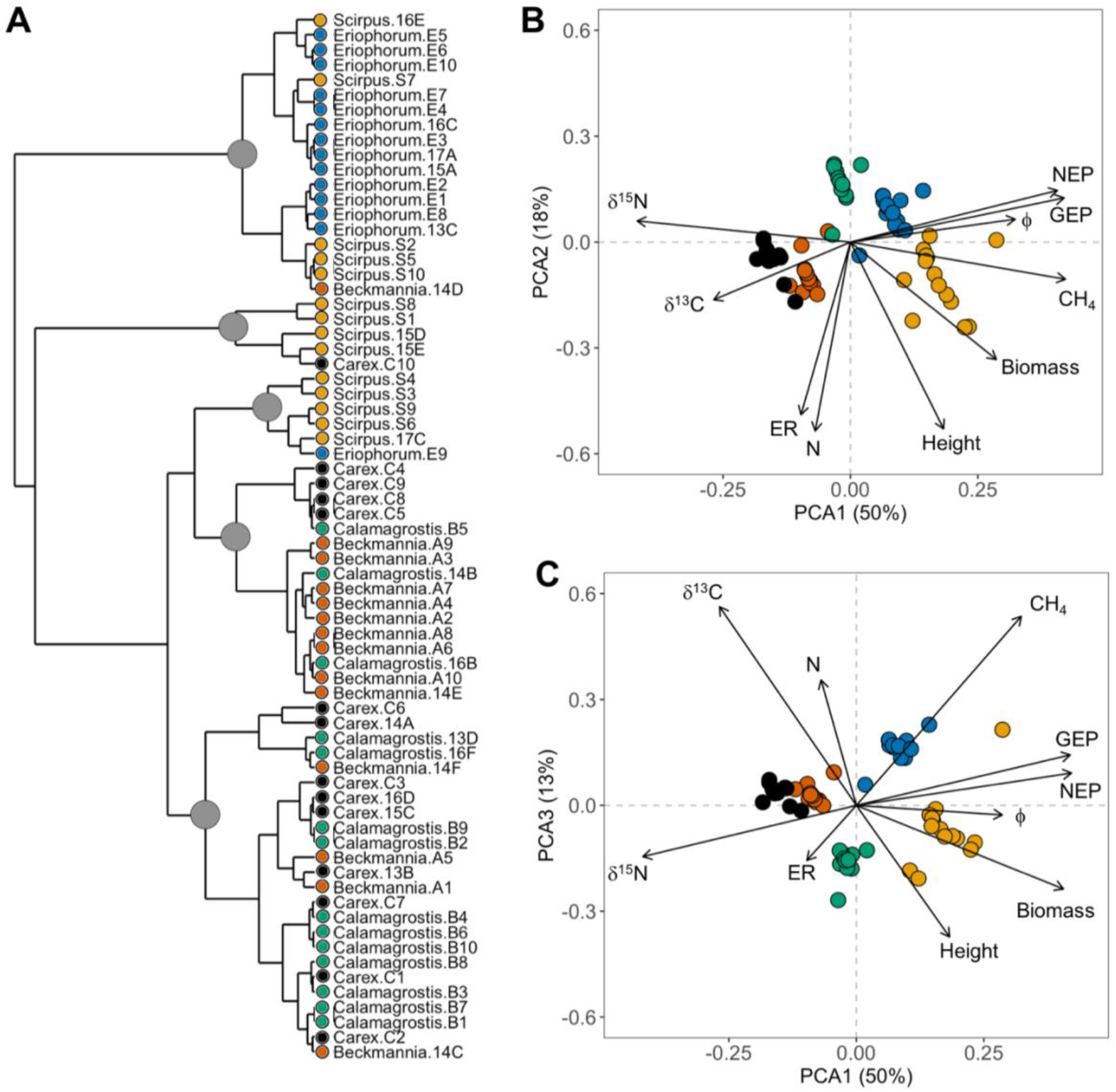
Functional variation among five graminoid species (n = 14 per species) based on net ecosystem CO_2_ production (NEP), gross primary productivity (GPP), ecosystem respiration (ER), methane (CH_4_,), above-ground biomass, plant height, quantum efficiency (φ), leaf nitrogen (N), leaf carbon and nitrogen stable isotope ratios (δ^13^C, δ^15^N). Groups were identified using (A) Ward’s agglomerative clustering and (B-C) principal components analysis (PCA). Significant clusters are indicated at the nodes (filled circles). PCA biplots show the strength of associations (% variance) between plant traits and species distributions along (B) PC1 (50%) and PC2 (18%); and (C) PC1 and PC3 (13%).

**Table 2:**
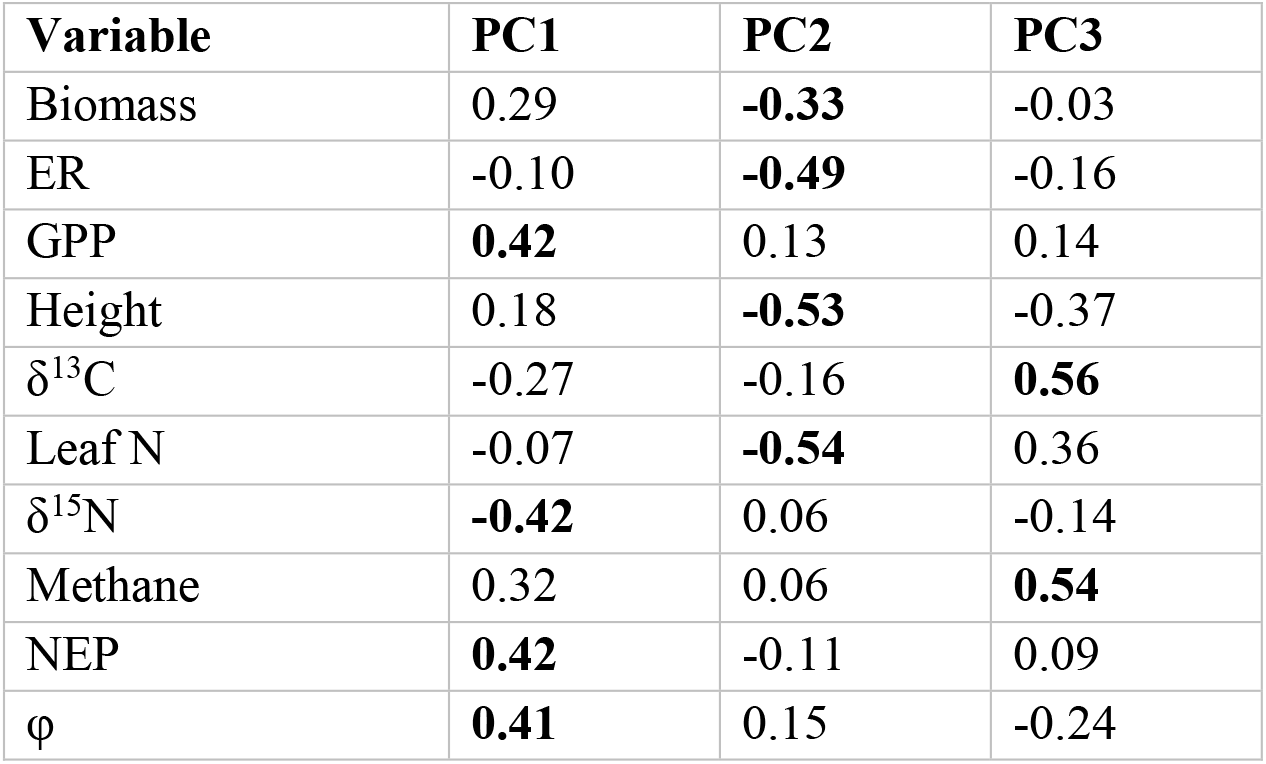
Eigenvector loadings for the first three principal components analysis (PCA) axes. Trait loadings describe the strength of associations (% variance explained) between the distribution of five graminoid species and ten plant descriptors: net ecosystem CO_2_ production (NEP), gross primary productivity (GPP), ecosystem respiration (ER), methane (CH_4_), above-ground biomass, plant height, quantum efficiency (φ), leaf carbon isotope composition (δ^13^C), and leaf nitrogen isotope composition (δ^15^N). Bold text indicates the corresponding PC axis for each variable. Data are from plot means (n = 14 per species).

| Variable | PC1 | PC2 | PC3 |
| --- | --- | --- | --- |
| Biomass | 0.29 | **-0.33** | -0.03 |
| ER | -0.10 | **-0.49** | -0.16 |
| GPP | **0.42** | 0.13 | 0.14 |
| Height | 0.18 | **-0.53** | -0.37 |
| $\delta^{13}\text{C}$ | -0.27 | -0.16 | **0.56** |
| Leaf N | -0.07 | **-0.54** | 0.36 |
| $\delta^{15}\text{N}$ | **-0.42** | 0.06 | -0.14 |
| Methane | 0.32 | 0.06 | **0.54** |
| NEP | **0.42** | -0.11 | 0.09 |
| $\phi$ | **0.41** | 0.15 | -0.24 |

Species did not clearly group together by plant family or environmental preference in cluster or principal components analyses (Figure 3). However, in both analyses *Scirpus* and *Eriophorum* formed one functional grouping while *Beckmannia*, *Calamagrostis* and *Carex* formed a second. These two groups differed in PC1 (F = 214.2, p < 0.0001) and PC3 (F = 8.341, p = 0.0052) and associated variables: NEP (F = 207, p < 0.0001), GEP (F = 124.5, p < 0.0001), δ^15^N (F = 1734, p < 0.0001), and φ (F = 91.81, p < 0.0001) for PC1 and CH_4_ (F = 129.4, p < 0.0001) for PC3 (Table 2). Although the two groups did not differ in PC2 (F = 0.434, p = 0.512) or associated variables of ER (F = 0.367, p = 0.547), leaf N (F = 0.037, p = 0.848), or plant height (F = 3.296, p = 0.0739), they differed in aboveground biomass (F = 32.67, p < 0.0001). The group with *Scirpus* and *Eriophorum* had larger rates of photosynthetic CO_2_ exchange and CH_4_ emissions, larger aboveground biomass, and leaves that were more light-use efficient (larger φ) and depleted in δ^15^N.

## Discussion

We assessed functional variation among five peatland graminoid species in order to better understand functional diversity in this important plant group. We tested hypotheses that CO_2_ and CH_4_ fluxes would vary due to differences in traits related to plant size and leaf physiology. Testing these hypotheses in a restored site presented a unique condition, as early ecosystem establishment provided a situation where the five species were growing in close association on similar substrate and hydrological conditions enabling more direct comparison among species. In support of our hypotheses, species differentially varied in CO_2_ and CH_4_ fluxes, with the sedge *Scirpus cyperinus* displaying the largest CO_2_ and CH_4_ fluxes followed by *Eriophorum vaginatum*. *Scirpus cyperinus* is common to freshwater wetlands, minerotrophic fens, and disturbed/agricultural areas in eastern North America from Georgia to Labrador (Ball et al., 2002) and can spontaneously re-establish (i.e., without active restoration techniques) in post-harvested peatlands (Cobbaert et al., 2004; Mahmood & Strack, 2011). *Eriophorum vaginatum* is abundant across northern North America in minerotrophic and ombrotrophic peatlands (e.g., fens and bogs) (Ball et al., 2002) and can also spontaneously re-establish on bare peat (Marinier et al., 2004). Moreover, *E. vaginatum* acts as a companion species to facilitate the growth and establishment of other peat-forming species, especially *Sphagnum* mosses, during early stages of restoration and peatland succession (Marinier et al., 2004; Tuittila et al., 2000). *Beckmannia syzigachne, Calamagrostis canadensis*, and *Carex canescens* had similar gas exchange rates (Figure 1). These three species are widely distributed across North American wetlands (Ball et al., 2002; Clark & Kellogg, 2007) and are less able to colonize on bare peat without restoration techniques (e.g., introducing donor material from natural areas, rewetting, straw mulch) (Cobbaert et al., 2004; González & Rochefort, 2019).

### Plant anatomical and physiological drivers of gas fluxes

Differences in gas fluxes among species were driven by variation in plant size and leaf physiological traits related to resource-use. Specifically, plants with larger gas flux rates had accumulated more above-ground biomass, were more light-use efficient (larger φ) and had relatively depleted leaf nitrogen stable isotope composition (δ^15^N). In fact, the strongest overall predictor of CO_2_ and CH_4_ fluxes was leaf δ^15^N. Variation in leaf δ^15^N is due to soil processes that influence patterns of soil δ^15^N such as mineralization, immobilization and the relative abundance of different soil N sources (e.g., organic, mineral). When growing in similar soil conditions, foliar δ^15^N can reflect differences in plant N uptake strategies (Goud & Sparks, 2018). For example, depleted (i.e., more negative) leaf δ^15^N can indicate root uptake of soil N sources that are more depleted in δ^15^N, such as nitrate (NO3^-^) relative to ammonium (NH4^+^), or N from deep within the soil profile (Craine et al., 2015). In this study, plants with relatively depleted foliar δ^15^N had larger CO_2_ and CH_4_ fluxes. These were predominantly *Scirpus* and *Eriophorum*, which were also the largest plants overall in terms of aboveground biomass and height. Depleted foliar δ^15^N in these species may reflect deeper roots that can tap into unexploited pools of soil nutrients and water, conferring growth advantages (Albano et al., 2021). Additionally, deeper soils and rooting zones have the potential for larger CH_4_ production; as such, the positive correlation between foliar δ^15^N and CH_4_ fluxes may be reflecting more plant-mediated release of CH_4_ from deeper in the soil profile than from smaller shallow-rooted species (Noyce et al., 2014; Strack et al., 2017).

Leaf N varied among species but did not correlate with either CO_2_ or CH_4_ fluxes. Many factors influence leaf N, including variation in soil N, leaf lifespan, life history, and rates of plant resource acquisition. Although leaf N can correlate with photosynthetic rates and biomass accumulation (McJannet et al., 1995; Walker et al., 2014; Wright et al., 2004), this is not always the case as much of a leaf’s N can be allocated to non-photosynthetic proteins as well as structural and defensive compounds (Ghimire et al., 2017). Additionally, many plants accumulate leaf N under high-irradiance or water-limited environments to economize water use during photosynthesis (Schrodt et al., 2015; Wright et al., 2003). Here, all three sedges had similar leaf N while the largest and smallest N contents were in grasses *Beckmannia syzigachne* and *Calamagrostis canadensis*, respectively. This interspecific variation is likely due to differences in leaf lifespan and phenology, rather than instantaneous carbon fluxes. For example, *Beckmannia syzigachne* is an annual, which can have larger leaf N relative to co-occurring or closely related perennial species (Garnier & Vancaeyzeele, 1994).

CO_2_ and CH_4_ fluxes were also strongly predicted by quantum yield (φ) such that gas fluxes increased with increasing φ, following similar patterns as biomass and foliar δ^15^N. As a measure of photosynthetic light-use efficiency, φ reflects the efficiency with which absorbed light is ultimately converted into fixed carbon (Monteith, 1977). Given that photosynthesis depends on the quantity and quality of light absorption, it has long been recognized that more light-use efficient plants can achieve larger rates of CO_2_ gas exchange, growth, and resource acquisition (Körner, 1982), and our results are no exception. Interestingly, φ also positively correlated with CH_4_ fluxes. It is likely that the larger CO_2_ exchange associated with more light-use efficient plants provides more photosynthetic root exudates that stimulate belowground CH_4_ production (Lai et al., 2014; Ström et al., 2005).

In addition to light absorption, leaf-level carbon gain is driven by the average difference in leaf internal and air external CO_2_ concentrations (*ci/ca*). Leaf carbon isotope composition (δ^13^C) is proportional to *ci/ca*, providing an integrated measure of the balance between metabolic demand and supply of CO_2_ via diffusion through the leaf boundary layer and stomata. In general, more depleted (negative) δ^13^C values indicate faster carbon metabolism or abundant CO_2_ supply (e.g., high stomatal conductance) (Farquhar et al., 1989; Goud et al., 2019). As expected, δ^13^C correlated with CO_2_ exchange, although this relationship was mainly driven by the smallest GPP and most enriched δ^13^C in *Carex canescens*. When variation in foliar δ^13^C is driven by stomatal conductance, enriched δ^13^C indicates stomatal closure which co-limits CO_2_ and water vapor diffusion. The combination of enriched δ^13^C, low φ, and small size in *Carex canescens* could indicate an ecological strategy to conserve water resources at the expense of carbon gain (Angert et al., 2009; Goud et al., 2019). On the other hand, *Scirpus* had the largest CO_2_ and CH_4_ flux rates and was also the largest, most light-use efficient, and had relatively fast carbon metabolism (largest φ, depleted δ^13^C), indicating a strategy to maximize carbon gain across a range of light and water availabilities.

Positive relationships between plant size and carbon fluxes have been reported for peatland and tundra graminoid species due to variation in aboveground biomass (Kao-Kniffin et al., 2010; Laine et al., 2012) and leaf area (Goud et al., 2017; Street et al., 2007). Variation in plant size among co-occurring plant species can be related to light competition. For example, sedges in riparian fens displayed strong competition for light based on height and aboveground biomass (Kotowski et al., 2006). In support of this, the sedges in this study varied widely from each other in gas flux rates and traits (except leaf N). Although grasses differed in height, they had similar aboveground biomass, likely reflecting differences in leaf area (Goud et al., 2017).

### Graminoid functional groupings

We hypothesized that plant functional variation would be more related to taxonomy than environmental preferences, with grasses and sedges forming distinct groups. Although we identified discrete groupings, they were not defined by taxonomy or environment. Indeed, based on multivariate analyses, no single species could be completely differentiated from neighboring species. Rather, we identified two functional groups defined by continuous plant traits and carbon fluxes. One group had larger rates of photosynthetic CO_2_ exchange and CH_4_ emissions, larger aboveground biomass, leaves that were depleted in δ^15^N and more light-use efficient, while the second group displayed the opposite suite of traits. These trait combinations are consistent with established ecological strategies related to growth rate and leaf economics, representing plants with either a strategy to grow quickly and invest in resource capture or to prioritize structural investment and resource conservation (Dìaz et al., 2015; Goud et al., 2019; Wright et al., 2004).

Currently, one of the most widely used peatland carbon models is the Holocene Peatland Model (HPM; Frolking et al., 2010) and its derivatives (e.g., Tuittila et al., 2012; Quillet et al., 2015). HPM distinguishes between grasses and sedges, and further subdivides sedges into minerotrophic and ombrotrophic plant functional types. NPP values defined for these graminoid PFTs are considerably smaller than what we report here. For example, by converting fluxes into common units, grasses in HPM (e.g., *Calamagrostis stricta*) are defined by NPP ranging from 210 – 850 gC m^-2^ yr^-1^ compared to a range of 1600 – 1800 gC m^-2^ yr^-1^ found here and in recent ecological studies (Rupp et al., 2019). Similarly, NPP prescribed for minerotrophic (e.g., *Carex canescens*) and ombrotrophic (e.g., *Eriophorum vaginatum*) PFTs in HPM are far smaller than what we observed: 210 – 1130 gC m^-2^ yr^-1^ and 30 – 190 gC m^-2^ yr^-1^, respectively, compared to 1350 – 4100 gC m^-2^ yr^-1^ and 2950 gC m^-2^ yr^-1^. Other peatland carbon models that use a single graminoid PFT (e.g., Chaudhary et al., 2017; Wania et al., 2009; Wu & Blodau 2013) also report NPP values much smaller than reported here (e.g., 90 – 150 gC m^-2^ yr^-1^).

Values for productivity and other functional attributes of PFTs are based on average environmental conditions of a defined site, rather than the total range of ecophysiological possibilities. For example, graminoids classified as ‘ombrotrophic’ are assigned smaller NPP values than ‘minerotrophic’, due to environmental constraints of ombrotrophic conditions (e.g., acidic, anoxic soils) despite their potential for a wide range of productivities across different peatland types. Indeed, the sedge *Eriophorum vaginatum* is usually classified as ombrotrophic despite its prevalence in minerotrophic peatlands, including early-successional and restored sites such as in this study. This does not mean that subdividing the graminoid PFT based on discrete environmental affinities is invalid. Rather, it’s possible that the range of productivity and plant function is more related to site-specific conditions rather than inherent properties of species and plant groups *per se* (Messier et al., 2010, 2016). In other words, intraspecific trait variability or phenotypic plasticity driven by local environmental conditions may have large impacts on how we define and apply PFTs (Adler et al., 2018; Henn et al., 2018; Westerband et al., 2021). A promising area of future research would be to evaluate drivers of intraspecific trait variability within functional types to constrain PFT delimitations.

### Applications for empirical studies and modeling

We did not find strong evidence to support grouping different graminoid species into one plant functional type. We also did not definitively find support for grouping species by taxonomy or reported environmental preferences. We do, however, think that characterising graminoids into functional groups based on morphological and physiological attributes, such as plant size and physiological efficiency that represent identifiable plant strategies is a promising way forward and may be less subject to biases introduced by PFTs (Dìaz et al., 2015). In addition, characterizing species more by their site-specific environmental conditions than assumed habitat preferences may account for intraspecific variation and improve functional designations in both empirical and modeling studies (Violle et al., 2012).

One limitation of this study is the limited number of species. Although the five species in this study differed considerably from existing graminoid PFT designations in terms of functional variation and magnitude of carbon fluxes, we acknowledge that they represent only a subset of possible species and environmental conditions across the diversity of peatland graminoids. Future studies are warranted that consider additional grass and sedge species and other graminoids such as rushes (e.g., *Juncus*), which are prevalent in peatlands but vastly under-represented in modeling and empirical studies. Synthesizing existing and new field data with publicly available trait databases (e.g., TRY) could provide critical insights to improve the representation of graminoids in ecological studies and modeling efforts.

Improved models that include peatlands are needed to couple regional and global Earth system models to accurately describe vegetation change and accompanying feedbacks to climate change. Moreover, modifying current models for natural peatlands to account for restored peatlands is crucial to evaluate current and future carbon storage during peatland management (Wania et al., 2009). Although there are efforts to incorporate peatlands into such models, the oversimplification of vegetation remains a key challenge. Given the limitations of discrete PFTs, an emerging approach is to apply continuous plant traits either to inform and update PFT classifications or to replace PFTs with traits altogether (VanBodegom et al., 2011). Indeed, replacing PFTs with continuous traits is the direction that many systems are moving towards; however, this is a particular challenge for peatlands. Trait-based models are currently designed for vascular plants with stomata, but a major contributor to peatland vegetative biomass and carbon exchange lies in bryophytes, especially *Sphagnum* peat mosses, that lack vascular tissue and stomata (Laine et al., 2012). Established trait protocols and trait databases for bryophytes are still in the beginning stages, but we are hopeful that their development will allow for the identification and application of traits common to bryophytes and vascular plants to better represent vegetation in empirical studies and modeling of current and future carbon storage.

## Acknowledgements

We are grateful for field assistance from Anoop Deol, Daniel Luckhurst-Cartier and Scott Macdonald. Sun Gro Horticulture provided site access and logistical support. This work was funded by an NSERC Canada Research Chair to MS and an NSERC Collaborative Research and Development Grant (grant number: 437463) supported by the Canadian Sphagnum Peat Moss Association and its members to MS and IBS. The authors state that they have no conflicts of interest to declare.

